# A Bayesian Multi-Task Approach for Detecting Global Microbiome Associations

**DOI:** 10.1101/2020.01.08.897538

**Authors:** Farhad Hatami, Emma Beamish, Rachael Rigby, Frank Dondelinger

## Abstract

**Motivation:** The human gut microbiome has been shown to be associated with a variety of human diseases, including cancer, metabolic conditions and inflammatory bowel disease. Current statistical techniques for microbiome association studies are limited by relying on measures of ecological distance, or only allowing for the detection of associations with individual bacterial species, rather than the whole microbiome.

**Results:** In this work, we develop a novel Bayesian multi-task approach for detecting global microbiome associations. Our method is not dependent on a choice of distance measure, and is able to incorporate phylogenetic information about microbial species. We apply our method to simulated data and show that it allows for consistent estimation of global microbiome effects. Additionally, we investigate the performance of the model on two real-world microbiome studies: a study of microbiome-metabolome associations in inflammatory bowel disease (Beamish, 2017), and a study of associations between diet and the gut microbiome in mice (Turnbaugh *et al*., 2009). We show that we can use the method to reliably detect associations in real-world datasets with varying numbers of samples and covariates.

**Availability:** Our method is implemented using the R interface to the Stan Hamiltonian Monte Carlo sampler. Software for running our methods is available at https://github.com/FrankD/MicrobiomeGlobalAssociations.

## 1 Introduction

Recent years have seen an explosion in the amount of genomic sequencing data collected on a variety of organisms. One area of particular interest is the nascent field of metagenomics. Metagenomics is concerned with the study of genetic material that can be found in samples from diverse environments ranging from soil and water to body cavities (Wooley *et al*., 2010). The collection of bacterial genetic material identified in a sample is called the microbiome. For human health, the human gut microbiome has been shown to be associated with a variety of conditions, including colorectal cancer (Ahn *et al*., 2013), metabolic diseases (Le Chatelier *et al*., 2013) and inflammatory bowel disease (Halfvarson *et al*., 2017).

The recent glut of microbiome studies has led to increased interest in developing robust and efficient statistical methods for analysing this type of data. A typical microbiome study will first sequence the bacterial genetic material contained in a sample, and will then use the genetic information to identify and quantify the organisms that the material originated from. In the case of the microbiome, the latter is most commonly done by using the 16S ribosomal RNA genes, which allow for the identification of bacterial species that colonize the human gut. Each fragment of 16S genetic material in the sample can then be classified by comparing it to known reference genomes, and species can be differentiated from each other by looking at genetic similarity. The result will be a count matrix counting the number of times genetic material from each species was detected.

Previous approaches to analysing microbiome data can be classified in two main categories: distance-based methods and regression-based methods. In distance-based methods we define a distance metric between observed microbiome samples. Common measures include Jaccard (dis)similarity, Bray-Curtis dissimilarity and the UniFrac distance (Lozupone and Knight, 2005). The latter has the advantage that it takes into account the phylogeny, or genetic relatedness, of the species in the samples when calculating the distance; loosely speaking, samples with related species will be closer than samples with unrelated species. One can then use a statistical test such as the PERMANOVA to test whether the centroids and dispersion of samples within two or more groups are identical (see e.g. Chen *et al*., 2012).

Regression-based methods tend to treat the count data as following either a negative binomial (Zhang *et al*., 2017) or Dirichlet-multinomial distribution (Chen and Li, 2013). In the former, the effects of observed covariates on species are usually treated as independent, although variance parameters may be shared across species. In the latter, the covariance across species is defined by the Dirichlet-multinomial distribution, but the regression will still define individual parameters representing the association between each species and the observed covariate. Unlike distance-based methods, regression methods do not try to infer a global effect of a covariate on the microbiome. However, they are better able to deal with multiple continuous covariates, and are not sensitive to a choice of distance measure.

In our work, we combine the advantages of the regression and distance-based approaches. Our idea is to develop a Bayesian multi-task log-ratio regression model, where the multi-task nature of the model allows us to model multiple species as multiple outcomes. Unlike many other regression approaches for microbiome data, we do not model the species counts directly, but rather treat the data as proportional and apply the log-ratio transform (Aitchison, 1982) to obtain multivariate-Gaussian distributed pseudo-observations. This allows us to bring in the phylogenetic information via the covariance matrix. Global associations between the microbiome and the covariates are estimated using a shared hyper-prior on the species-specific effect estimates. We demonstrate the robustness and efficiency of our method using an extensive simulation study, and apply it to a case study in inflammatory bowel disease.

## 2 Methods

### 2.1 Log Ratio Model

Let us assume that we are collecting microbiome data from *N* people, and in each sample we are counting the occurrence of *S* species or taxons. We treat the microbiome data as compositional, i.e., even though we are observing count data, we only consider the proportion of each species. If we observe an *S*-dimensional count vector **z**_*i*_ for sample *i*, then the proportion of species *s* can be estimated as 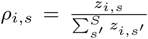. We can now transform the species proportions for species *s* ∈ {1, …, *S* − 1} by using the log-ratio transform (Aitchison, 1982):

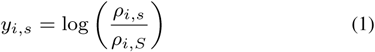

We assume that the **y**_*i*_ are approximately multivariate Gaussian distributed, and model them as:

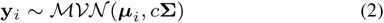

In our application, **Σ** is informed by the phylogenetic tree inferred from the sequencing data. Briefly, under the assumption of a Brownian motion evolutionary model, the covariance of two species is equivalent to the branch length from the root to the most recent common ancestor in the phyogenetic tree. If no phylogenetic tree is available, then we set **Σ** = **I**, the (*S* − 1) × (*S* − 1) identity matrix. The hyperparameter *c, c* > 0 is a scaling parameter controlling the desired fit to the data. For the data in this study we found that *c* = 1 led to satisfactory models that showed no evidence of over-or under-fitting.

### 2.2 Multi-task regression

Having established the log ratio model, we now want to formulate ***µ****_i_* as a linear expression with information sharing across the species (“tasks”, or response variables). Let:

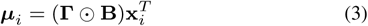

where **x**_*i*_ is a vector of *p* observed covariates for subject *i*, **B** is an (*S* − 1) × *p* matrix of positive linear coefficients and **Γ** is an (*S* − 1) × *p* matrix with entries in {−1, 1}. Estimation of global effects is achieved via a prior on the elements of ***β***_*j*_, the *S* − 1-dimensional vectors corresponding to the columns in **B**:

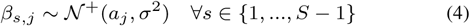

where *a*_*j*_ denotes the global effect of covariate *j* on all species. Here *𝒩*^+^ denotes the positive part of the Gaussian distribution truncated at zero. Note that the global effect only defines the prior for the strictly positive magnitude *β*_*s,j*_ of the species-specific effects. Placing a prior on the signed effect size would not have the desired effect; since the size of each biological sample is fixed, an increase in some species must lead to a decrease in other species, and so the global effect size under such an alternative model would not be reflective of the change across species.

We also need to define a distribution for the elements *γ*_*s,j*_ of **Γ**. The natural approach would be to treat them as binary variables following a Bernoulli distribution. However, this requires us to perform inference over discrete variables, which can be onerous, and does not lend itself well to gradient-based inference techniques. Instead, we opt for a Gumbel-Softmax approximation to the Bernoulli distribution (Jang *et al*., 2016; Maddison *et al*., 2016). The Gumbel-Softmax is a continuous relaxation of the categorical distribution. In general, for a categorical random variable *h* with class probabilities *π*_1_, *π*_2_, …, *π*_*K*_, we can represent *h* as:

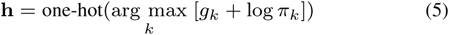

where one-hot denotes the one-hot encoding of a categorical variable as a *K*-dimensional vector (hence the change from *h* to **h**). The *g*_*i*_ are sampled from a Gumbel(0,1) distribution. Note that this representation is exactly equivalent to a categorical distribution; to obtain the continuous relaxation we need to replace the arg max by a softmax function, which gives:

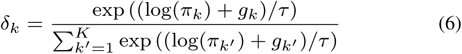

for *k* ∈ {1, …, *K*}, where ***δ*** ∈ Δ^*K*−1^, the (*K*−1)-dimensional simplex. Here *τ* is a tuning parameter that controls the smoothness, with *τ* = 0 reverting to the discrete case. In our case, *K* = 2, which means that eq. (6) simplifies to:

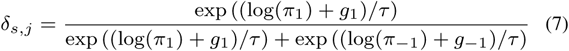

where *π*_−1_ = 1 − *π*_1_. We drop the subscript *k* as one value of *δ* fully defines the two-dimensional vector on the simplex, but introduce the subscript (*s, j*) to denote the value of *δ* for species *s* and covariate *j*. We use the subscripts 1 and −1 to denote Gumbel draws for *γ*_*s,j*_ = 1 and *γ*_*s,j*_ = −1 respectively. Since *δ*_*s,j*_ is either 0 or 1, we set:

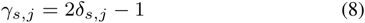

This has the desired property that if *δ*_*s,j*_ ≈ 1, *γ*_*s,j*_ ≈ 1 and if *δ*_*s,j*_ ≈ 0, *γ*_*s,j*_ ≈ −1.

We specify truncated Gaussian *𝒩*^+^(0, 1) priors for the global effects *a*_*j*_, which also serves to regularize the model when the sample size is smaller than the number of parameters, which is the case for almost all the models studied here. We further place a Beta prior on the parameter *π*_1_, where the hyperparameters can be used to encode any prior beliefs about the prevalence of positive or negative associations; for our study we set both equal to 1.

### 2.3 Inference

We are interested in performing fully Bayesian posterior inference over the parameters of this model. A convenient framework is Hamiltonian Monte Carlo (HMC), which allows us to take advantage of the differentiability of the Gumbel-Softmax to estimate gradients and speed up the Bayesian inference. We employ the Stan sampler to perform HMC sampling; for more details on the the Stan sampler and modelling language see Carpenter *et al*. (2017).

In microbiome research, it is common to encounter datasets consisting of large numbers of operational taxonomic units (OTUs). Despite the efficiency of HMC sampling, sampling hundreds or thousands of parameters *β*_*s,j*_ and *γ*_*s,j*_ is not currently feasible on a personal computer. We therefore propose an approximate estimation, as an alternative to the full log-ratio model, where we only estimate *β*_*s,j*_ as part of the sampling, and the *γ*_*s,j*_ parameters are estimated in a separate inference step. The approximate inference procedure is as follows:

1. Fit independent linear models 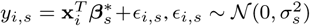 to the log-transformed microbiome counts for all species *s*. Note that 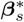 is distinct from the rows of **B** in the full model.

2. Set 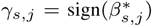 where the sign() operation returns −1 if 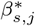 is negative and 1 otherwise.

We refer to the model with **Γ** inferred from the data as the full log-ratio model, and to the model with **Γ** fixed according to the two-step procedure above as the conditional model, since inference is conditional on the estimate of **Γ**. We can think of the conditional model as an approximation to the full joint model *P* (**Γ, B, *θ***|**Y, X**) = *P* (**B, *θ***|**Γ, Y, X**)*P* (**Γ**|**Y, X**), where ***θ*** collects all parameters not in **B** or **Γ**, and ***X*** and ***Y*** represent the log-ratio response and the design matrix with the observed covariates, respectively. Instead of estimating *P* (**Γ** | **Y, X**), which would require marginalising over all other parameters, we use 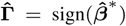 where 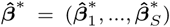 is the maximum likelihood estimate of the effect sizes in independent linear models of 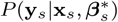 for each *s*. Inference is performed on the conditional model 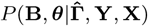, which is less computationally expensive.

## 3 Simulation Study

### 3.1 Setup

We perform a simulation study under different scenarios to test the performance of the full and conditional log-ratio models. In particular, we study how well the full model can predict the global effect (*a*_*j*_) and species-specific effect (*β*_*s,j*_) parameters under scenarios with different sample size *n*, variance parameter *σ*^2^, and number of species (or taxonomic units) *S* and compare the performance to that of the conditional model. In each experiment the variables that are not changing are set to *n* = 100, *σ*^2^ = 0.1 and *S* = 10. The global effect sizes in the simulation model are set fixed with *a*_*j*_ ∈ {0, 0.5, 1}.

For each unique setting of the simulation parameters, we generate 20 random datasets according to the following procedure. First, a random phylogenetic tree is generated using the ape R package (Paradis and Schliep, 2018). Based on this tree, the covariance matrix **Σ** under the Brownian motion evolutionary model is calculated. Next, we generate an *N* × *p* matrix of covariates **X** with *x*_*i,j*_ ∼ *𝒩* (0, 1). For each covariate *j* and species *s, γ*_*s,j*_ are generated from a binomial distribution with *p* = 0.5. Species-specific coefficients *β*_*s,j*_ are generated according to eq. (4). Then we simply calculate ***µ***_*i*_ according to eq. (3), and sample ***y****i* according to eq. (2). The *ρ*_*i,s*_ must sum to 1, and, for *s* ∈ {1, …, *S* − 1}, must be equal to *ρ*_*i,S*_ exp(*y*_*i,s*_) by eq. (1). To achieve this we set 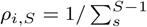 exp(*y*_*i,s*_). We can then calculate the *ρ*_*i,s*_ and simulate count data **z**_*i*_ from a multinomial distribution with parameters ***ρ***_*i*_, and number of trials equal to 100 *\* S*.

The bench-marking platform used for this study ran R-3.6.1 to generate datasets and invoke `rstan`. Models were fitted using the Stan software (Carpenter *et al*., 2017) version 2.19.1. A comparison between popular methods such as the PERMANOVA (generalized uniFrac method using the `GUniFrac` R package) (Chen and Chen, 2018), ANOSIM (non-parametric multivariate analysis using the `vegan` R package) (Clarke, 1993) and MiRKAT (microbiome regression-based kernel association test using the `MiRKAT` R package) (Zhao *et al*., 2015) has been done. The sampling was executed in parallel using 4 different chains each with 8000 iteration with a stopping criteria at convergence on 4 Intel Ivy Bridge cores each running at 1.15GHz speed and 12GB of RAM memory in total.

### 3.2 Results

In this section we investigate and compare the performance of:

- the full log-ratio model,
- the conditional log-ratio model which uses a linear marginal model for each species to determine the species-specific effect signs *γ*_*s,j*_ and then infers the size of the species-specific effects *β*_*s,j*_ and the global effect *a*_*j*_ conditional on the *γ*_*s,j*_,
- the “oracle conditional model”, where the *γ*_*s,j*_ parameters are set to the true values used for simulating the data.

Figure 1 shows the performance of the models for estimating the global effect size in three different experiments: changing the sample size, noise variance on the *β*_*s,j*_ parameters, and number of species. Dashed lines represent the true values of the global effect (*a*_*j*_) in the simulation model. It can be seen that increasing the number of samples *n* improves the accuracy of prediction and reduces the variance. On the other hand, increasing the variance parameter for the effect values and increasing the number of species (or taxonomic units) both make estimation of the global effect parameters harder. We see that there is not much difference between the model performance for the three log-ratio models, but as could be expected, the full log-ratio model produces less biased estimates compared to the conditional log-ratio model, with the oracle log-ratio model outperforming both.

**Fig. 1:**
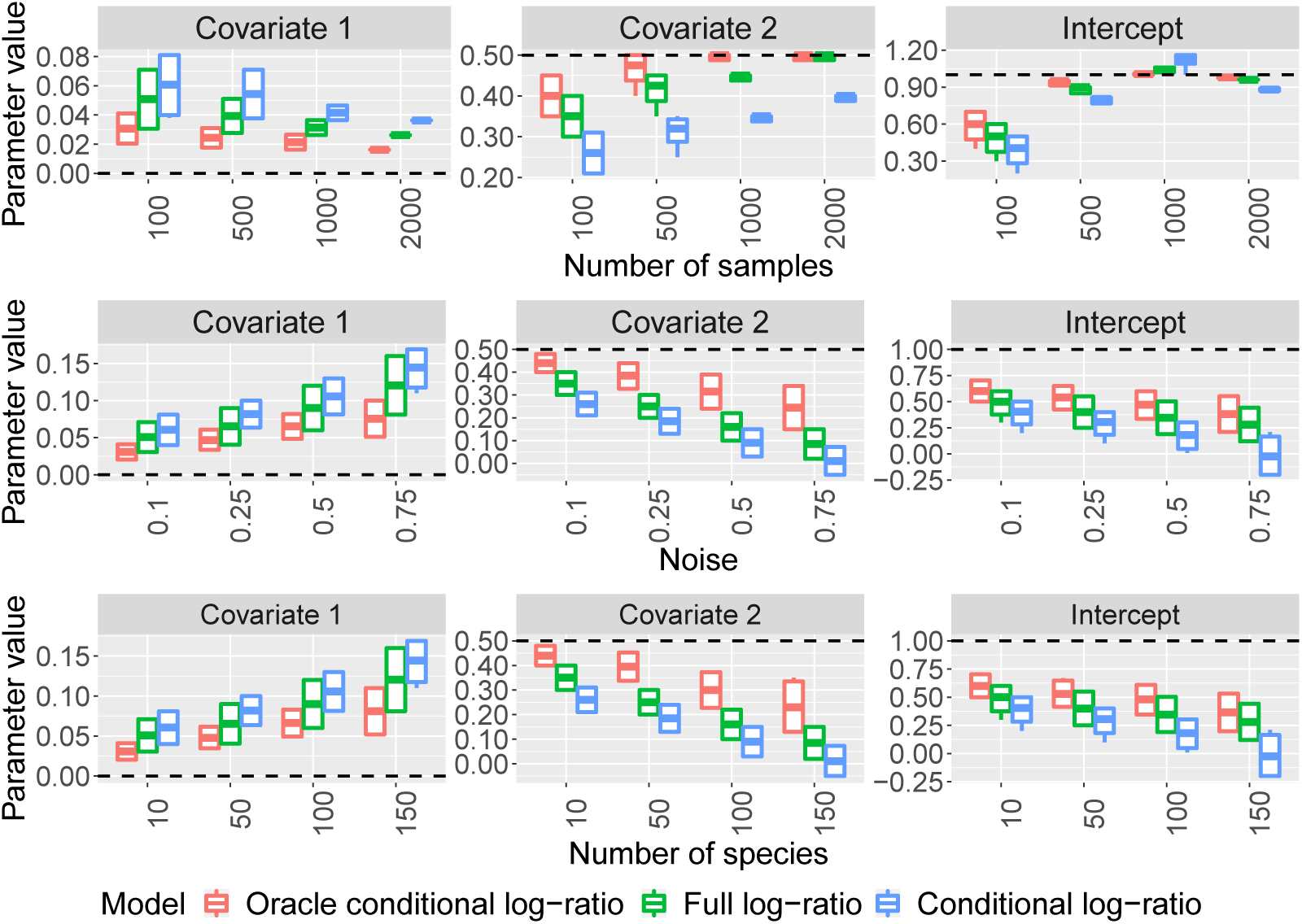
Comparison of the performance of the full, conditional and oracle conditional log-ratio models on simulated data when varying the sample size (top row), noise variance of the species-specific effects (middle row) and number of species or taxonomic units (bottom row). The linear simulation model consists of the intercept term and two covariates with global effect sizes 1, 0.5 and 0 respectively. The dashed horizontal lines show the size of the true effect, and the boxplots represent the posterior mean estimates obtained from 20 independent simulated datasets.

Figure 2 shows a comparison between the performance of the three models when predicting the species-specific effect size *β*_*s,j*_. We note that species-specific effects are predicted well across all three models. Looking at the both figures 1 and 2 we can see in most cases the full log-ratio model outperforms the conditional model.

**Fig. 2:**
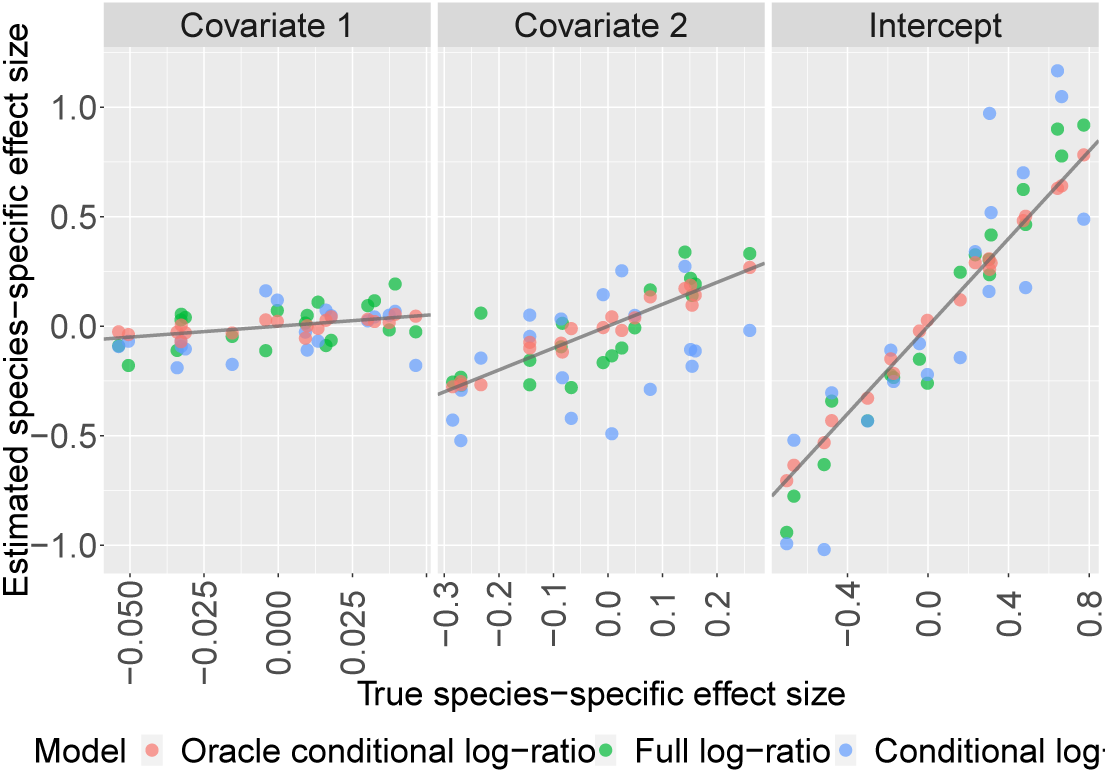
Comparison of the performance of the full, conditional and oracle conditional log-ratio models on simulated data when estimating the species-specific effect size (*β*_*s,j*_). Each point corresponds to the estimated effect size for one species in one model. The linear simulation model consists of the intercept term and two covariates with global effect sizes 1, 0.5 and 0 respectively. The solid lines represent *y* = *x* (note that the x-axes are on different scales).

Figure 4a shows a comparison between the *p*-values obtained using the PERMANOVA (generalized UniFrac), ANOSIM and MiRKAT tests performed on some simulated data when increasing the number of covariates. The PERMANOVA, ANOSIM and MiRKAT models quantify the effect of a covariate by the percent variability in microbial compositions and do not have a straightforward comparison with our log-ratio models beyond *p*-value. Figure 4a shows that when the number of covariates exceeds 30 then *p*-value> 0.05 and hence we reject the null hypothesis (which is the good estimation of the global effects). This is due to the fact that all these three distance-based methods run a significant test for each covariate separately.

**Fig. 3:**
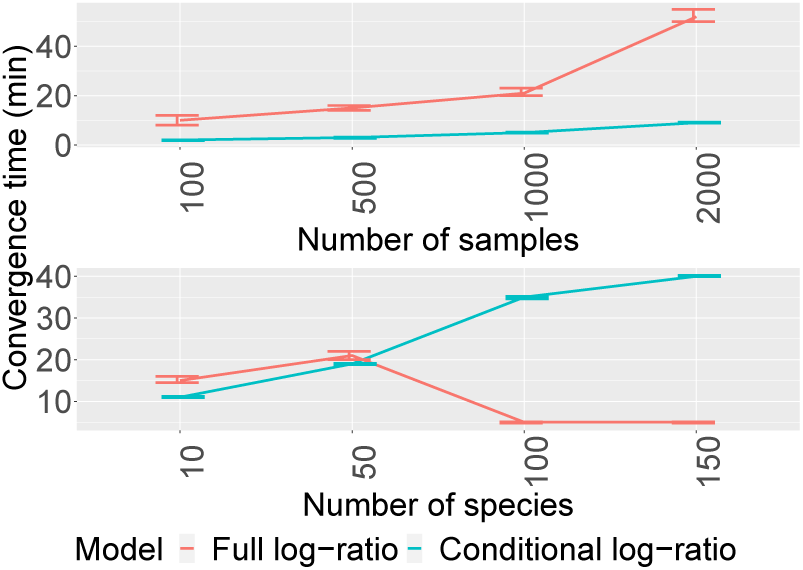
Comparison of the computational time to convergence of the full and conditional log-ratio models, on the simulated data when increasing sample size *n* and number of species (or taxonomic units) *S*. The error bars show the 95% confidence interval calculated over 20 independent simulated dataset.

**Fig. 4:**
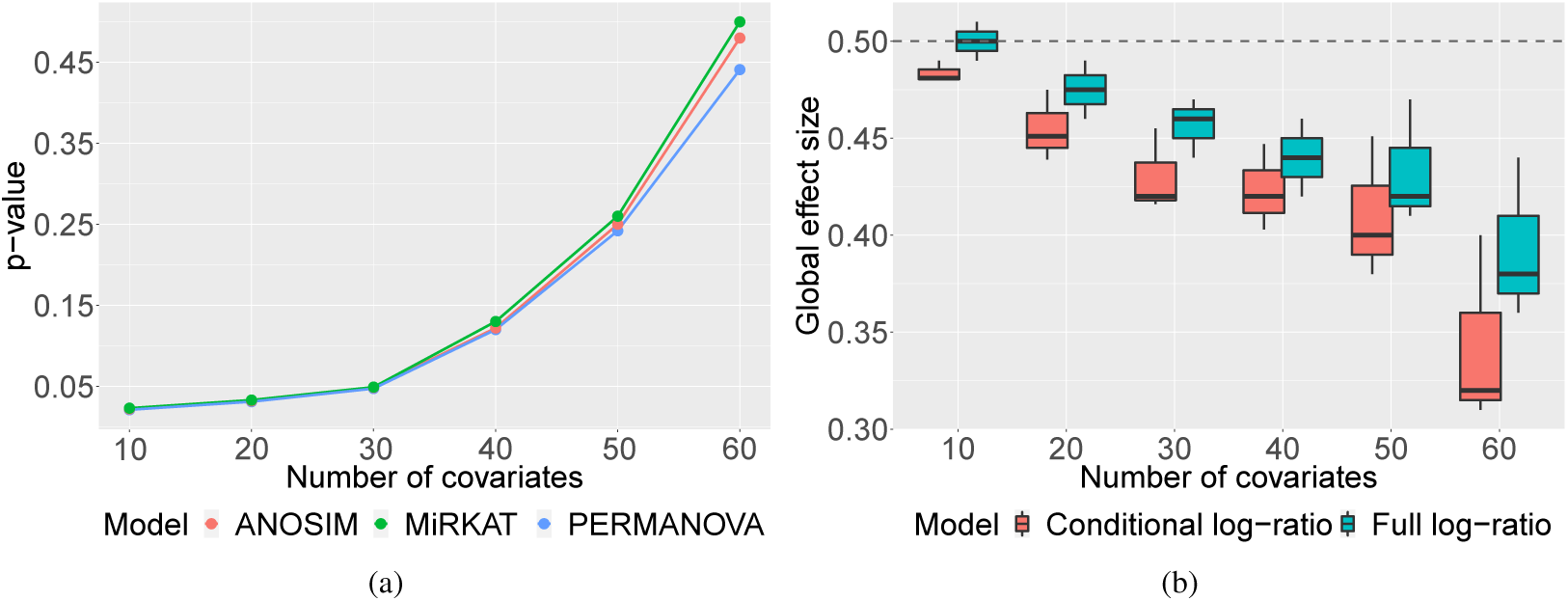
(a) Comparison of the performance of the ANOSIM, MiRKAT and PERMANOVA models on simulated data when increasing the number of covaritaes. (b) Comparison of the performance of the full and conditional models on simulated data when increasing the number of covaritaes. The y axis represents the estimated global effects with the dashed line representing the mean value.

The generalized UniFrac approach estimates a single effect parameter using a distance measure based on all species; by comparison, our method uses a hierarchical model to estimate both global effects and species-specific effect, making it more flexible.

Figure 4b shows a comparison between the performance of the full and conditional log-ratio models when increasing the number of covariates. We see that both models perform reasonably well for higher number of covariates. Note that while these results are not directly comparable with the p-values in Figure 4a, we notice that the estimated global effect size remains far from zero under our models, indicating a lower false negative rate.

We also investigate the computational efficacy of the full and conditional log-ratio models. Figure 3 shows the run time to convergence under each of the models. We see that the sampler for the conditional model takes longer for large *S*; this is a consequence of setting the *γ*_*s,j*_ parameters fixed which results in less efficient exploration of the sample space and longer convergence times.

Taken together, Figures 1, 2 and 3 indicate that for small sample sizes (*n* < 100) one could use either the full or conditional log-ratio models. For large sample sizes the conditional model could be deployed to improve computational efficiency, but there is a trade-off in terms of estimation accuracy. For large number of species *S* (*S* > 50), the full log-ratio model seems more efficient than the conditional model, perhaps because it can take advantage of Hamiltonian paths in the posterior landscape that are closed off when conditioning on **Γ**.

## 4 Real-World Applications

### 4.1 Detecting Global Microbiome-Metabolome Associations

Using metabolomics as a measure of health, or to predict response to pharmacologics is a very attractive prospect, as measuring metabolites in urine, for example, may provide a non-invasive and potentially inexpensive screening method that has potential for application to a plethora of pathologies.

We apply the full log-ratio model to a dataset collected as part of a pilot study into inflammatory bowel disease (IBD) (Beamish, 2017). IBD is an umbrella term used to describe two conditions (Crohn’s disease and ulcerative colitis) that are characterized by chronic inflammation of the gastrointestinal tract. 42 gut bacteria samples were obtained from patients undergoing colonoscopy at Lancashire Teaching Hospitals Trust (LTHTR) and University Hospitals of Morecambe Bay Trust (UHMBT), and metagenomic profiling was performed to obtain the bacterial species counts. The patients were additionally asked to provide a urine sample, and metabolite concentrations found in those samples were recorded. One objective of the study was to investigate associations between the gut microbiome and the urinary metabolite profiles.

After quality control, our cohort consists of *n* = 40 people (IBD cases and controls), for which we have measured a set of 3 clinical covariates: age, sex, and the hospital where the sample was collected. Note that we do not investigate any associations with IBD status, as number of cases was too small to make any reasonable inferences. Instead, we focus on detecting associations between the microbiome and individual metabolites. We consider the set of the *S* = 21 most abundant operational taxonomic units (OTUs or species) at the phylum level. As a result, our design matrix **X** is a 40 × 5 matrix consisting of the intercept, the metabolite level, age, sex and sample collection location. As the data is limited, we do not try to infer the variance parameter *σ*^2^, but keep it constant and equal to 0.1. Finally, we have a total of 261 metabolites.

Figure 5a shows the posterior distribution of estimated global effect size *a*_*j*_ for the metabolites with the largest posterior mean effect estimate. We see that out of these 21 metabolites, saccharopine, allantoin, vanillate and ornithine have the strongest association with the microbiome. Some metabolites show interesting bimodal patterns in their posterior effect distribution; this is particularly obvious for glycine. Such patterns could potentially be indicative of more than one mechanism that links the microbiome to metabolite levels.

**Fig. 5:**
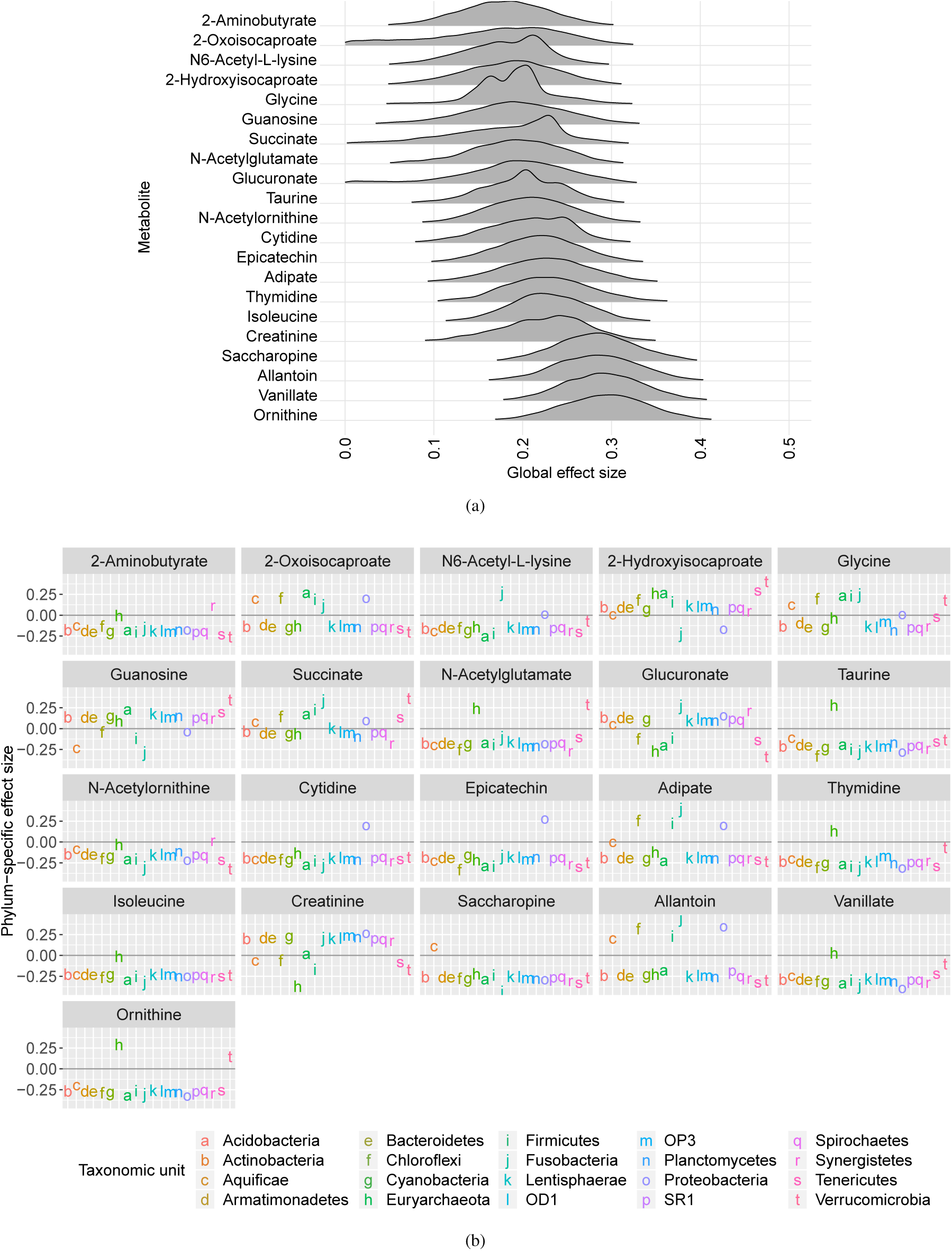
Application of the log-ratio model to a microbiome-metabolome association study (Beamish, 2017). (a) Density plots showing the posterior distribution of the global effect size associating each metabolite to changes in the microbiome. (b) Posterior mean estimates of the phylum-specific effect size for each metabolite across the different bacterial phyla.

Figure 5b shows the posterior mean estimate of the species-specific effect sizes *β*_*s,j*_, specifically the association of each phylum with each metabolite. We note that for the metabolites with the largest posterior effects, the majority of species-specific effects are strongly negative, indicating that these metabolites are associated with a decrease in most phylum counts, compensated by an increase in just a couple of phyla.

We note that our analysis identified several phyla-associated metabolites that have previously been shown to be linked to markers of intestinal health or nutrition, demonstrating potential impacts on health that are reflective of microbiota changes. First, our study suggests that adipate is strongly associated with changes in the microbiome. Adipate, the salts and esters of adipic acid, are increased in the urine of breast-fed vs. formula-fed neonates (Dessì *et al*., 2016), a now well-known essential factor in the establishment of a diverse, healthy microbiome. We observe effect sizes that are indicative of an association between adipate and an increase in Firmicutes, Proteobacteria and Bacteroidetes, the major components of a healthy adult intestinal flora. Therefore, clinical measurement of adipate may have potential as biomarkers for intestinal health.

Epicatechin is a polyphenolic flavonoid abundantly present in plants, cocoa and that offers protection against diabetes by its ability to mimic the effects of insulin and has strong antioxidant properties (Kanehisa and Goto, 2000; Samarghandian *et al*., 2017). The association we observed between epicatechin and proteobacteria abundance may prove useful in the prediction of pathologies associated with proteobacteria expansion, such as IBD or nonalcoholic fatty liver disease (Rizzatti *et al*., 2017). Of course, any potential mechanisms that underpin these associations will require further investigation but collectively this demonstrates the broad and significant potential for clinical application of our model. Several metabolites, including isoleucine, thymidine, ornithine and vanillate, were associated with an increase in cyanobacteria (Figure 5b). Cyanobacteria are unusual intestinal microflora (Ley *et al*., 2005), in that they perform oxygenetic photosynthesis and cyanobacteria-based supplements, such as Spirulina, regarded as functional foods, have been shown to support immunological function (Finamore *et al*., 2017). These associations may allow us to get a clearer picture of the functionality of the gut microbiome, by assessing the functions of these metabolites.

### 4.2 Estimating the Global Effect of Diet on the Gut Microbiome

It is well-known that diet has a profound effect on the composition of the gut microbiome (David *et al*., 2014; Singh *et al*., 2017). However it is difficult to quantify this effect using traditional microbiome analysis methods, as we either need to look at individual species, or rely on a pre-specified distance metric. In Turnbaugh *et al*. (2009), the authors took the latter approach to characterise the differences in microbiota in mice whose gut tracts were colonised by donor bacteria of human or mouse origin, and who received either a Western or low-fat plant-based (LF/PP) diet. The dataset is described in Table 1. The dataset consists of *n* = 444 fecal samples, obtained at various time points from 15 mice, and we use the bacterial abundance data for *S* = 160 taxonomic units at the genus level. Note that we did not have access to the information on sample times or sample source, and so were unable to take this aspect into account in our model.

**Table 1.**
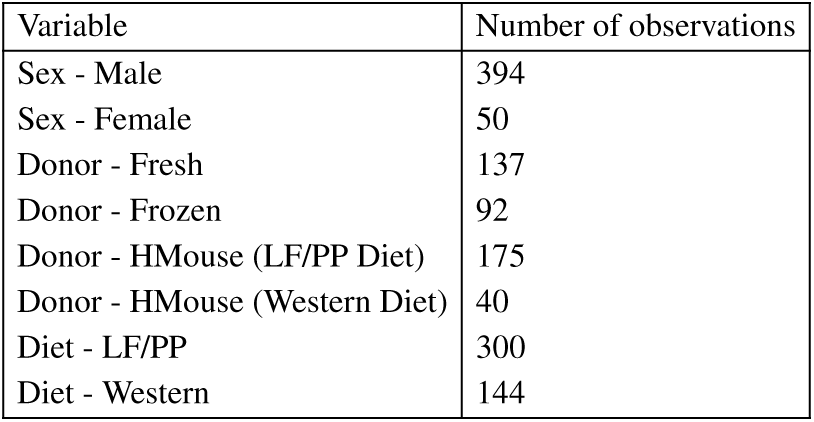
Dataset description for the gut microbiome study in Turnbaugh et al. (2009). Donor bacteria either originated from human microbiome samples (fresh or frozen), or from mice who had previously been implanted with (fresh) human samples (HMouse) and had received either the LF/PP or Western diet.

| Variable | Number of observations |
| --- | --- |
| Sex - Male | 394 |
| Sex - Female | 50 |
| Donor - Fresh | 137 |
| Donor - Frozen | 92 |
| Donor - HMouse (LF/PP Diet) | 175 |
| Donor - HMouse (Western Diet) | 40 |
| Diet - LF/PP | 300 |
| Diet - Western | 144 |

Using the UniFrac distance measure, Turnbaugh *et al*. (2009) found that recipient mice consuming a Western diet showed a similar microbiome composition that was distinct from mice on the LF/PP diet, regardless of donor. We applied the full log-ratio model to the dataset to determine whether this was reflected in the global microbiome effect of diet on the microbiome, and to study the individual (genus-specific) effects. We include sex, donor type and diet as covariates. The reference taxonomic unit for the log-ratio transform was the Dorea genus, although investigation of alternative reference taxonomic units showed little change in the results (see supplementary information).

The results of our study are shown in Figure 6. We observe that overall there is a significant (non-zero) effect of diet, and a strong effect of using the frozen samples. This is concordant with the findings in (Turnbaugh *et al*., 2009). We also note that receiving a bacterial implant from a donor mouse on the LF/PP diet seems to have a stronger effect than receiving an implant from a donor mouse on the Western diet, compared to having a human donor. Ruminococcaceae Incerta Sedis is strongly associated with the change in diet, a finding which is backed up by the literature (Clarke *et al*., 2013). Similarly, Eubacteria colonization of the gut has been shown to be affected by diet (Simmering *et al*., 2002). Finally, we see that Enterobacteria is associated with the change in diet; previous studies in rats have shown that a high-fat diet results in considerably more propionate and acetate producing species, including Enterobacteria (Singh *et al*., 2017). Interestingly, this bacterial genus also shows a strong association with implantation by frozen donor bacterial samples.

**Fig. 6:**
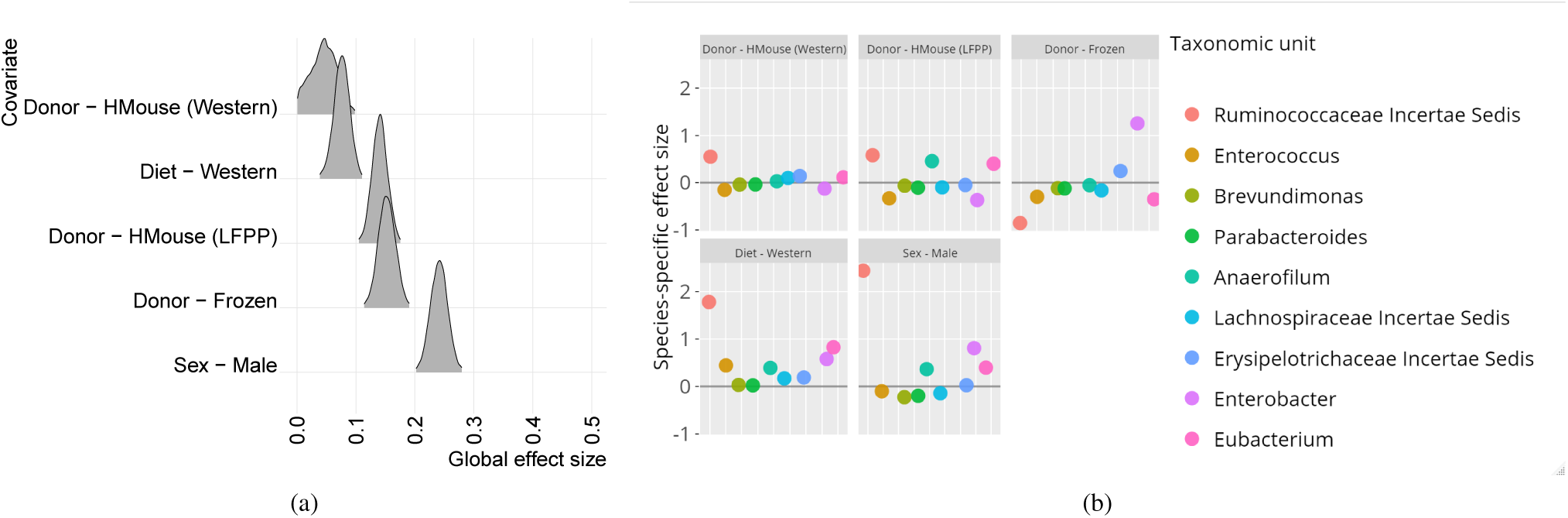
Application of the log-ratio model to a gut microbiome study (Turnbaugh *et al*., 2009). The reference taxonomic unit for the log-ratio transformation is Dorea. (a) Density plots showing the posterior distribution of the global effect size associating each covariate to changes in the microbiome. Note that the reference category for diet is LF/PP, meaning that the global effect is the change with respect to that diet. Similarly the reference category for bacterial donor is fresh (human), and the reference category for sex is female. (b) Posterior mean estimates of the genus-specific effect size for each metabolite across the different bacterial genera. We display here the 10 OTUs with the highest effect size for the association with diet.

## 5 Conclusion

We have developed a Bayesian multi-task regression model for detecting global microbiome associations, and have shown that the model produces consistent estimates in a simulation study. We then applied the model to investigate microbiome-metabolome associations in a pilot study of inflammatory bowel disease patients and global associations with diet in a study of mouse gut microbiomes. The model is superior to existing approaches in that it allows for integrating phylogenetic information, it estimates global as well as species-specific effects, and it does not rely on a specific choice of ecological distance measure.

There are some limitations to our study. The IBD dataset used in this paper only consists of a small number of samples, which reduced our ability to extend the inference to a larger number of species. For a larger sample size, the full log-ratio model could also be applied to taxonomic units at the ‘order’ or ‘genus’ level, as our simulation study showed that inference for up to 150 species is feasible without major modifications. However, we have seen that for large numbers of samples (*n ≥* 500), the conditional log-ratio model can be deployed as it showed comparable prediction accuracy to the full model at potentially greater computational efficiency.

An interesting avenue for future research is whether our model can scale to larger numbers of species *S* and sample sizes *n*. We note that time to convergence seems to increase polynomially with n; we would therefore not expect to be able to scale to tens of thousands of samples, and a different inference method would need to be employed. One option is variational inference, which has been successfully used as an alternative to full Bayesian inference in other settings (Hensman *et al*., 2013).

More crucially, we would like the method to scale to larger numbers of species. Currently, our results indicate that HMC inference in the full model converges faster as *S* increases. Our hypothesis is that for small *S*, the gradient estimate for *a*_*j*_ is unreliable as it depends on the estimates of *β*_*s,j*_. For large *S*, we have more information to estimate the gradients, and thus experience fewer rejections during HMC sampling. However, estimation of the parameters will be poor unless *n* also increases. In future work, we will investigate regularization approaches to estimate sparse *β*_*s,j*_ parameters in situations with small sample sizes.

## Acknowledgements

We thank Dr Albert Davies for collection of patient samples.

